# When and where to find the tiny creatures: environmental predictors of the abundance of a miniaturized frog species from Northeast Brazil

**DOI:** 10.1101/2025.07.22.666076

**Authors:** Laiana Carla de Moura, Igor Joventino Roberto, Murilo Guimarães, Flávia Ribeiro Bezerra, Kássio Castro Araújo, Ednilza Maranhão dos Santos

## Abstract

Amphibians are the most vulnerable terrestrial vertebrates to environmental changes, primarily due to their permeable skin and dependence on water for reproduction. In miniaturized frogs, sensitivity to changes in ecological factors related to humidity may be particularly critical due to their high surface-to-volume ratio, which makes them more susceptible to desiccation. To test this hypothesis, we evaluated the influence of environmental variables on the abundance of calling males of *Adelophryne nordestina*, a small miniaturized frog with direct development inhabiting the Atlantic Forest leaf litter in northeastern Brazil. We surveyed calling males of *A. nordestina* over nine months across four transects, each containing 25 survey points, totaling 100 study sites. We employed a Negative Binomial N-Mixture Model to estimate the abundance of calling males and assessed the effect of environmental parameters. Our results indicate that estimated mean detection probability was 0.03, and mean abundance among sites was 85.90. Precipitation, temperature, leaf litter depth, and canopy cover significantly influenced the number of calling males in the study area. We suggest that, as a terrestrial, miniaturized, and direct-developing species, changes in environmental parameters affecting microhabitat humidity impact its reproductive activity. Our findings contribute to the understanding of miniaturized frog ecology and the potential impacts of environmental disturbances on this group.

## Introduction

Environmental conditions are well known to affect species distribution and composition assemblages, especially in ectothermic species, which depend directly on specific microclimate to survive (Isaak et al. 2017). Among vertebrate ectothermics, amphibians have permeable skin and non-amniotic eggs (Wells 2007). Because of this, they are primarily influenced by environmental factors like temperature, rainfall, humidity, and solar irradiation (Steelman and Dorcas 2009; Cook et al. 2011; Ospina et al., 2013; Alton & Franklin, 2017; Bonnefond et al. 2020; Ceron et al. 2020). Amphibians are one of the most threatened groups among vertebrates, in which recurrent population declines have been reported in the last decades (IUCN 2022). Habitat loss, fragmentation, diseases, contaminants, alien species introduction, overexploitation, and global climate changes are the primary causes of this decline (Beebee & Griffiths 2005; Collins 2010; Green et al. 2020). Hence, comprehending the relationship between amphibians and environmental factors is paramount to mitigating population declines and securing species persistence (Pellet & Schmidt 2005).

Among the amphibians, miniaturized frogs can be especially susceptible to environmental impacts. Animal miniaturization involves a series of changes in morphology, ecology, natural history, and behavior (Hanken & Wake 1993; Yeah 2002; Rittmeyer et al. 2012). Miniaturized frogs usually inhabit the leaf litter of humid forests, have direct development, and produce a small number of eggs per spawning (Rittmeyer et al. 2012). In addition, they are more vulnerable to desiccation due to their high surface-to-volume ratio; therefore, a preference for more humid climates is favorable for their development (Tracy et al. 2010). It is hypothesized that miniaturization has allowed certain species to exploit microenvironments inaccessible to larger anurans, including smaller prey and shelter from predators (Kraus et al. 2011; Scherz et al. 2019).

*Adelophryne* Hoogmoed and Lescure, 1984 is among the miniaturized frogs from humid forests of South America. The genus is composed of 12 species with a mean body size of 2-3 cm and direct development (Hoogmoed & Lescure 1984; Hoogmoed et al. 1994; MacCulloch et al. 2008; Santana et al. 2012; Taucce et al. 2020; Lourenço-de-Moraes et al. 2012, Lourenço-de-Moraes et al. 2014, Lourenço-de-Moraes et al. 2018, Lourenço-de-Moraes et al. 2021). Except for the threatened species, *A. maranguapensis* Hoogmoed, Borges, and Cascon, 1994 (see Cassiano-Lima et al. 2011, 2014, 2020; Oliveira et al. 2022; Araújo et al. 2023) and *A. baturitensis* (Araújo et al. 2023), ecological studies are scant on these tiny frogs. Their small size, cryptic coloration, limited distribution, and secretive habits in nature are some of the reasons for the lack of ecological studies on *Adelophryne* (Fouquet et al. 2012) and other miniaturized frogs, making them challenging species to monitor in nature. In addition, most of them were described in the last decade such as *A. nordestina* (Lourenço-de-Moraes et al. 2021), which is distributed in the northern Atlantic Forest, in the Pernambuco Endemism Center, Brazil’s most fragmented and threatened sub-biogeographical region (Ribeiro et al. 2009). As a terrestrial species with direct development and high sensitivity to desiccation - traits associated with its miniaturized nature - we hypothesize that *A. nordestina* is particularly vulnerable to microhabitat environmental changes. In this way, here we assess the effects of precipitation, humidity, temperature, leaf litter, and canopy cover on the species’ reproductive activity. Our goal is to shed light on niche dimensions of miniaturized frogs from tropical forests and their susceptibility to environmental changes. Given the challenges posed by climate change and other threats to amphibians, this knowledge is crucial for guiding the group’s monitoring and conservation planning.

## Material and methods

### Study area

The present study was carried out in the Parque Natural Municipal Professor João Vasconcelos Sobrinho (hereafter, PNJVS), municipality of Caruaru, Pernambuco state, Northeast Brazil (-8.35530, -36.02868). The area is a conservation unit of 349 ha located in the “Brejo dos Cavalos”, an altitudinal Atlantic Rainforest enclave surrounded by Caatinga environments in the Brazilian semiarid (Tabarelli & Santos 2004; Andrade-Lima 2007). The PNJVS (Fig. 1) comprises large areas of forest in regeneration that had been used for agriculture crops and livestock. The more isolated regions have preserved forest remnants (SUDER/SES 2018).

**Figure 1.**
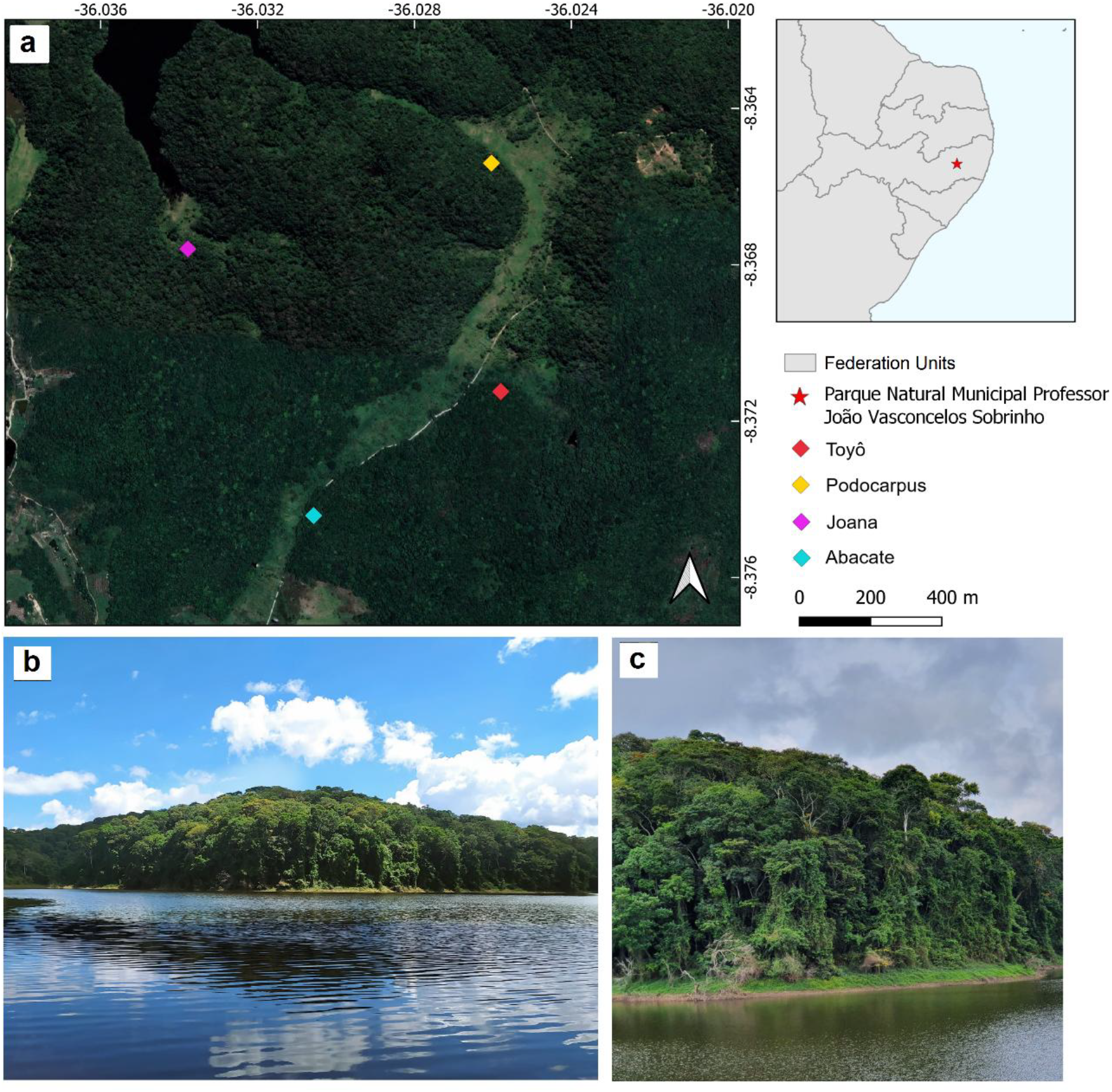
Study area showing the transects where individuals of *A. nordestina* were sampled (a). Typical environments in the Parque Natural Municipal Professor João Vasconcelos Sobrinho, Pernambuco state, northeast Brazil (b, c). The map was created by QGIS 3.28 (https://qgis.org). Photographs by LCM

### Sampling

We counted calling male individuals of *A. nordestina* in the PNJVS between 07:00 and 22:00 h in four secondary rainforest transects 250m long, each separated by at least 0.5 km (Fig. 1 and 2). All four transects were situated near small streams and were named after the local communities in their respective areas: Abacate, Podocarpus, Velha Joana, and Toyô. Transects were sampled monthly, for three days, from January to September 2022, totaling 27 sampling days. To estimate the number of calling males of *A. nordestina* each transect was divided into 25 sites 10 m apart from each other (a distance empirically determined to prevent recounting the same individual), and recorded the number of individuals calling at that site for approximately one minute, following the methodology described by Dorcas et al. (2009). The same researcher (LCM) conducted the counting to minimize bias, and an additional independent researcher (FRB) assisted in confirming the number of vocalizing frogs in most field surveys. Hence, here we consider each site as an independent sampling unit, totaling 25 sites per transect, 100 in total. To validate our call identifications, we recorded samples of the advertisement calls and compared them with the species’ described advertisement call characteristics, finding no differences.

**Figure 2.**
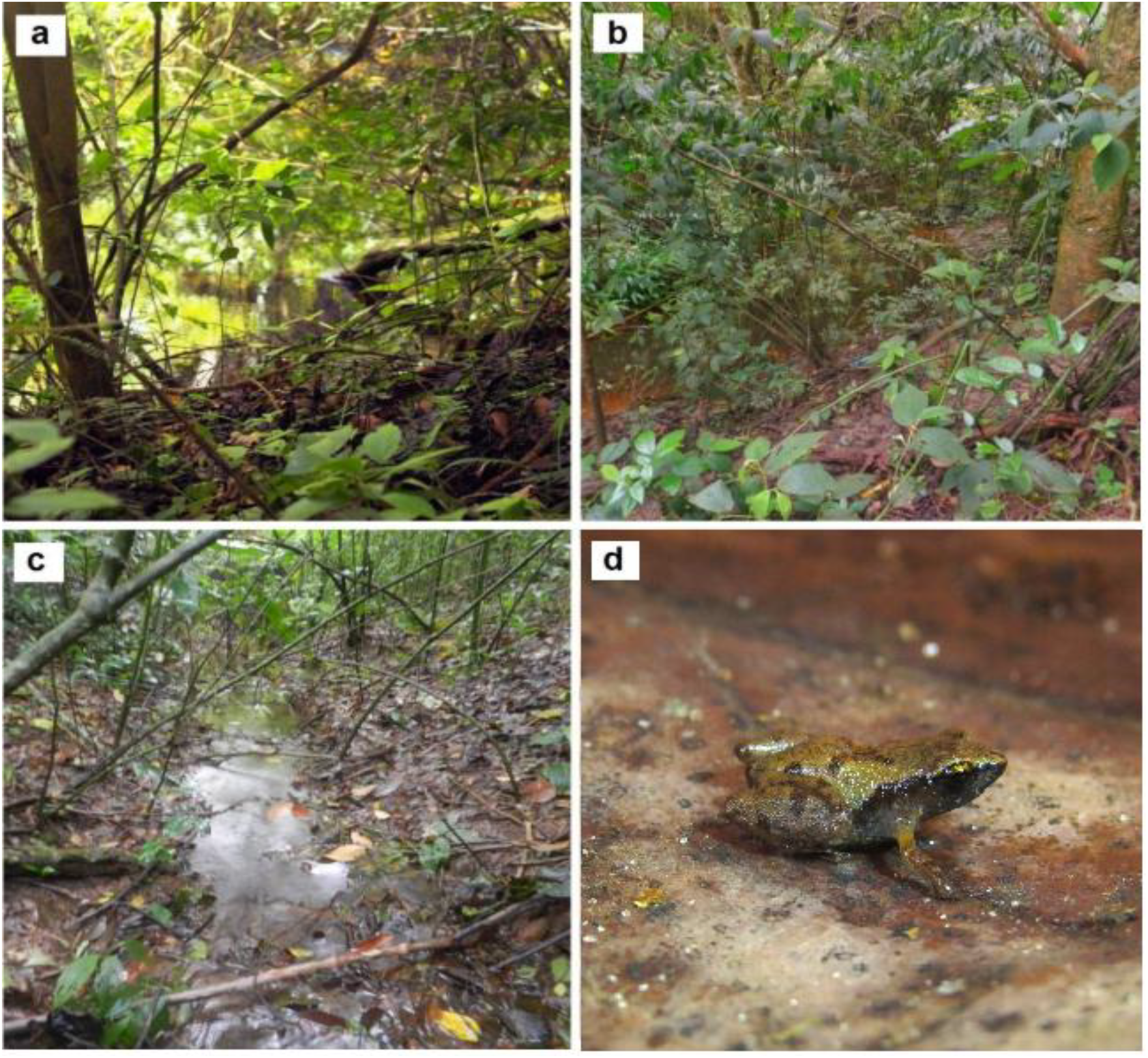
Forest habitats and small streams in sampled habitats of *Adelophryne nordestina* (d) in the Parque Natural Municipal Professor João Vasconcelos Sobrinho, Pernambuco state, northeast Brazil. Photographs by Anna Mello (a) L. Moura (b, c and d)

### Environmental variables

We included the following numeric temporal predictors and their quadratic terms, as additive effects: leaf litter, temperature, humidity, accumulated rainfall from the last 15 days, and accumulated rainfall from the last day. In addition, we included time, representing the study occasions to accommodate any temporal variability not captured by our temporal predictors. As spatial fixed predictors, we included the percentage of canopy openness and its quadratic term (numeric), and habitat (factor, with four levels representing the four transects we surveyed).

For the leaf litter depth, we selected 25 points uniformly distributed along each transect measured using a metric ruler with subdivisions of 1 mm. Likewise, we measured temperature and humidity using a thermometer reading Incoterm 7666. For precipitation in the previous 15 and 1 day before the surveys, we used the database available by the Agência Pernambucana de Águas e Clima (APAC, http://old.apac.pe.gov.br/meteorologia/monitoramento-pluvio.php#). We measured canopy openness percentage through digital photos taken by a Nikon D50 digital camera with a Nikon dx 18-105mm fisheye 67-58mm hemispherical lens. We used the Gap Light Analyzer 2.0 software to analyze the photos of each transect according to Frazer, Canham, and Lertzman (1999).

### Data analyses

All numerical predictors were z-transformed to have zero mean and one standard deviation. We tested numerical predictors for multicollinearity using correlation tests and the variance inflation factor (VIF). We excluded one predictor of a given pair whenever a high correlation (> 0.70; Dormann et al. 2007) was detected.

We obtained Maximum Likelihood estimates using closed-population N-mixture models within *unmarked* R package (Fiske & Chandler 2011) using the *pcount* function. The closed N-mixture model assumes that the population is demographically and geographically closed during the study period, abundance is random and independent among sites, observers do not double count individuals, and heterogeneity is fully modeled (Kéry & Schaub 2011). The N-Mixture model is a hierarchical model that provides estimates of abundance while accounting for sampling error, and the detection probability. The model requires counts *y*_*i,j*_ for a set of *i* sites and *j* visits, and such counts arise from two distinct processes, the ecological process and the observation process. The spatial variation of local abundance at each site *i* for a group of sites is driven by a count distribution, such as the Poisson, with mean *λ*. The observed counts *y*_*i,j*_ given *N*_*i*_ at site *i* and visit *j* are described by a Binomial distribution with mean *N*_*i*_ and detection probability, *p*_*i,j*_ (Kéry & Schaub 2011).

Given that our data set was composed of nine months we split it into two parts based on the breeding season of the species and climate seasonality. The first part, called season 1, spread from January to April and represented the pre-breeding season, characterized by higher temperatures than season 2, from May to September, which was characterized by the breeding season and lower temperatures. With the data set divided into two seasons, we performed a season-stratified analysis using a stacked data set (Kéry & Royle 2021). Stacking data is a solution when seasons are roughly independent and there is not enough temporal replication to implement an open-population model (Kéry & Royle 2020). Considering all predictors included in the analysis, our initial model for abundance and detection probability were, respectively: log(*λ*) = α_0_+ α_1_*canopy.open + α_2_*I(canopy.open^2) + α_3_*habitat + α_4_*season and logit(p_*i,j*_) = β_0_ + β_1_temperature + β_2*_I(temperature^2) + β_3_rainfall_1 + β_4_I(rainfall_1^2) + β_5_rainfall_15 + β_6_I(rainfall_15^2) + β_7_*humidity + β_8_*I(humidity^2) +β_9_leaflitter + β_10_*I(leaflitter^2) + β_11_time.

We compared three types of mixtures using our data set: Poisson, Negative Binomial, and Zero-inflated Poisson. To select the best mixture, we built a full model, containing all selected predictors, and assessed model fit using a parametric bootstrap goodness of fit test with *Nmix*.*gof*.*test* function in *AICcmodavg* R package (Mazerolle 2013), and visual inspection of residual plots.

After selecting the most appropriate mixture, we started from the full model and performed a backward stepwise variable selection (Zuur et al. 2009) to find the best predictors explaining our parameters of interest. We started with the detection probability submodel, and then followed to the abundance submodel excluding terms based on the hypothesis testing procedure, dropping the least significant terms, one by one, until only significant terms remained.

## Results

We observed the individuals of *A. nordestina* inhabiting the surroundings of small streams about 1m wide, in forested environments and were in calling activity during all months from January to September 2022 in the PNJVS. The individuals called continuously from 07:00 am to 10:00 pm. The monthly mean of calling males was 58.66 ± 25.8, in which the highest abundance (96 individuals) was found in May and the lowest (22 individuals) in August. The months with the highest abundance of calling males were April, May, and July, and those with the lowest abundance were March, August, and February. The average temperature during transects was 22.6ºC, ranging from 19.73ºC (August) to 26.03ºC (March), and the humidity was 75% (March) to 94.25%, (June), with an average of 88.6%. The leaf litter depth mean value was 3.98 cm, the mean accumulated rainfall in the previous 15 days was 37.40 mm^3^, and on the previous day of the survey was 2.15mm^3^. The canopy openness had a mean of 14.33% among the 100 surveyed sites, varying between 4.66% and 35.74%.

Among our covariates, temperature and the accumulated rainfall from the last 15 days were correlated (r = 0.73), and we kept the temperature in our analysis. Similarly, humidity and rainfall from the last 24 hours were also correlated (r = 0.89), and we kept rainfall. The Negative Binomial mixture presented the best fit to our data with the full model showing the smallest overdispersion (c-hat = 1.26) and the smallest AIC value. The Negative Binomial is a mixture that includes another parameter to be estimated, the dispersion parameter, related to the variance of mean abundance, estimated as 0.66 (CI 0.58 – 0.73).

According to our best model, the estimated mean detection probability was 0.03 (CI 0.01 – 0.04), and the mean abundance among sites was 85.90 (CI 54.40 – 135.70). Our best model included temperature, leaflitter, time, and the quadratic effect of rainfall as important predictors of detection probability. Abundance differed according to the quadratic effect of canopy openness and among the four habitats (table 1). Detection probability increased linearly with temperature and leaf litter whereas it showed a non-linear relationship with rainfall (Table 1, Fig. 3). Detectability also varied among visits (Table 1, Fig. 3). Mean abundance varied among sites and habitats, from 2 to 194 individuals (Table 1, Fig. 4, Fig.S1). Canopy openness presented a non-linear relationship with abundance (Table 1, Fig. 5).

**Table 1.** Models results based on the best Negative Binomial N-mixture model for the abundance and detection probability submodels for male adults *Adelophryne nordestina*. The intercept for the abundance submodel represents habitat ‘abacate’, while for the detection submodel represents time 1 (first survey).

| Abundance: | Estimate | SE | z | P(>\|z\|) |
| --- | --- | --- | --- | --- |
| (Intercept) | 4.45 | 0.28 | 16.02 | 9.88e-58 |
| canopy.open | 0.19 | 0.10 | 1.94 | 5.30e-02 |
| I(canopy.open^2) | -0.11 | 0.05 | -2.21 | 2.73e-02 |
| habitatjoana | -0.73 | 0.21 | -3.55 | 3.88e-04 |
| habitatpodocarpus | -1.60 | 0.24 | -6.83 | 8.42e-12 |
| habitattoyo | -0.25 | 0.19 | -1.29 | 1.96e-01 |
| Detection: | Estimate | SE | z | P(>\|z\|) |
| (Intercept) | -3.68 | 0.30 | -12.49 | 8.02e-36 |
| temperature | 0.17 | 0.06 | 2.96 | 3.08e-03 |
| rainfall | 0.64 | 0.09 | 7.05 | 1.73e-12 |
| I(rainfall^2) | -0.25 | 0.06 | -4.46 | 8.06e-06 |
| leaf litter | 0.09 | 0.04 | 2.08 | 3.78e-02 |
| time2 | -0.70 | 0.13 | -5.30 | 1.15e-07 |
| time3 | -0.59 | 0.13 | -4.57 | 4.91e-06 |
| time4 | -0.62 | 0.13 | -4.68 | 2.83e-06 |
| time5 | -0.61 | 0.17 | -3.49 | 4.84e-04 |

**Figure 3.**
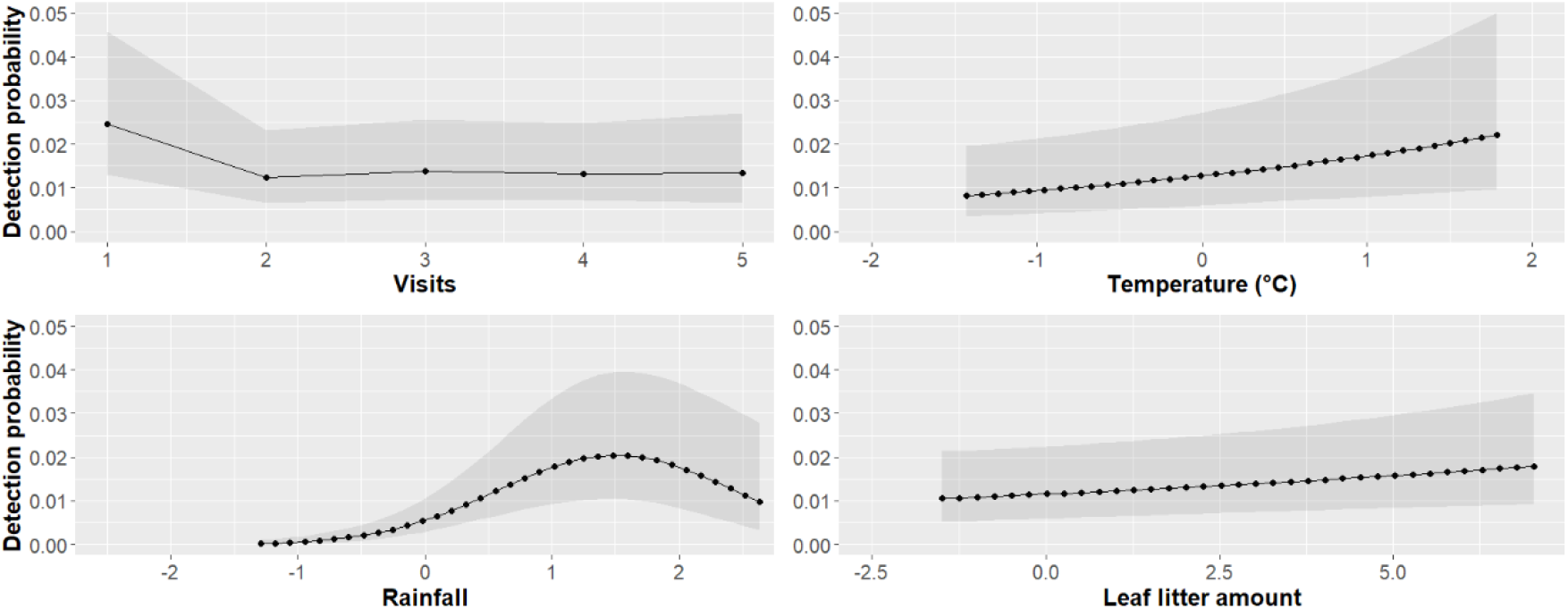
Scaled predictor effects on detection probability of *Adelophryne nordestina* adult males. The shaded area represents the 95% confidence interval

**Figure 4.**
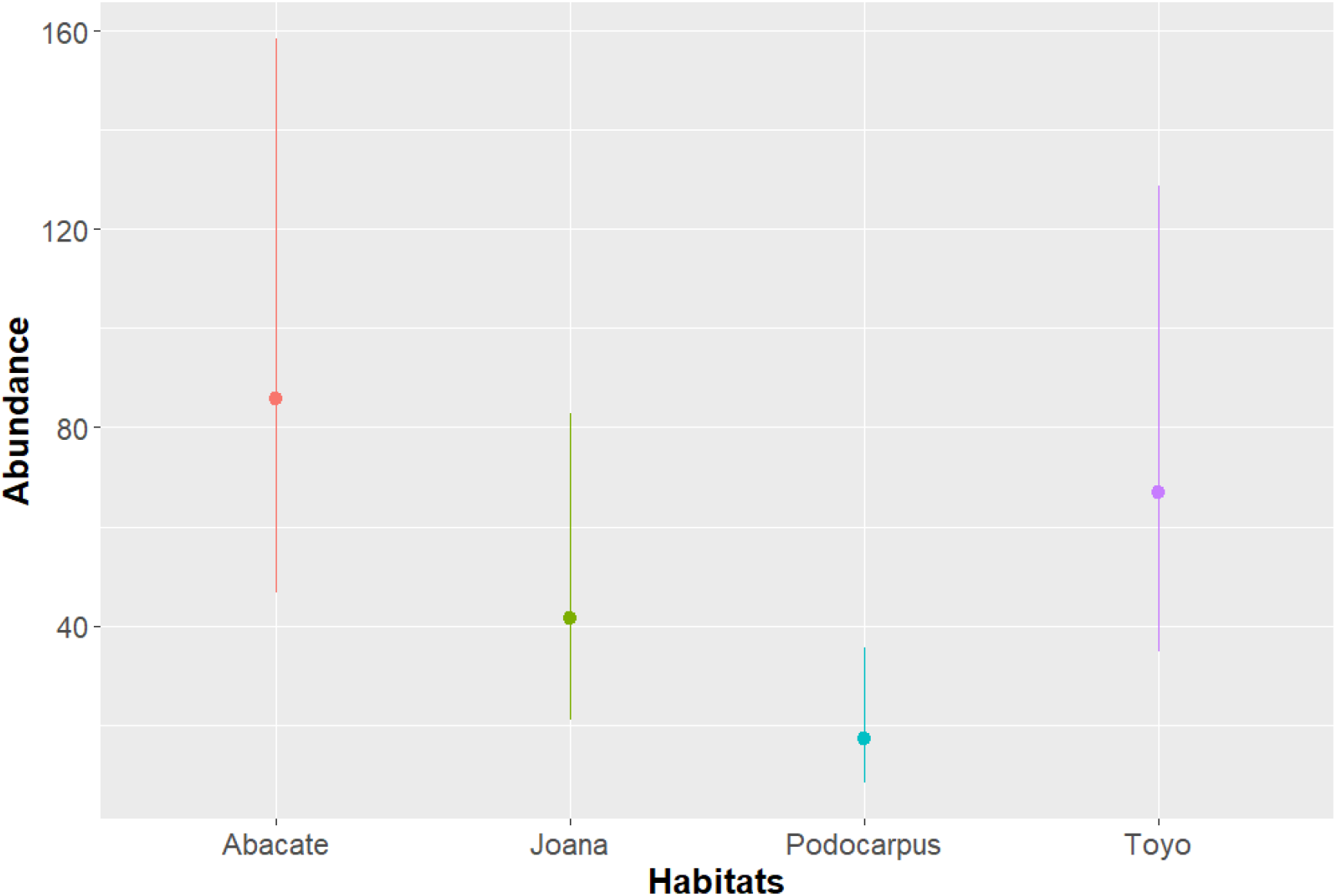
Mean abundance estimates and 95% confidence limits of *Adelophryne nordestina* adult males among habitats during the study period

**Figure 5.**
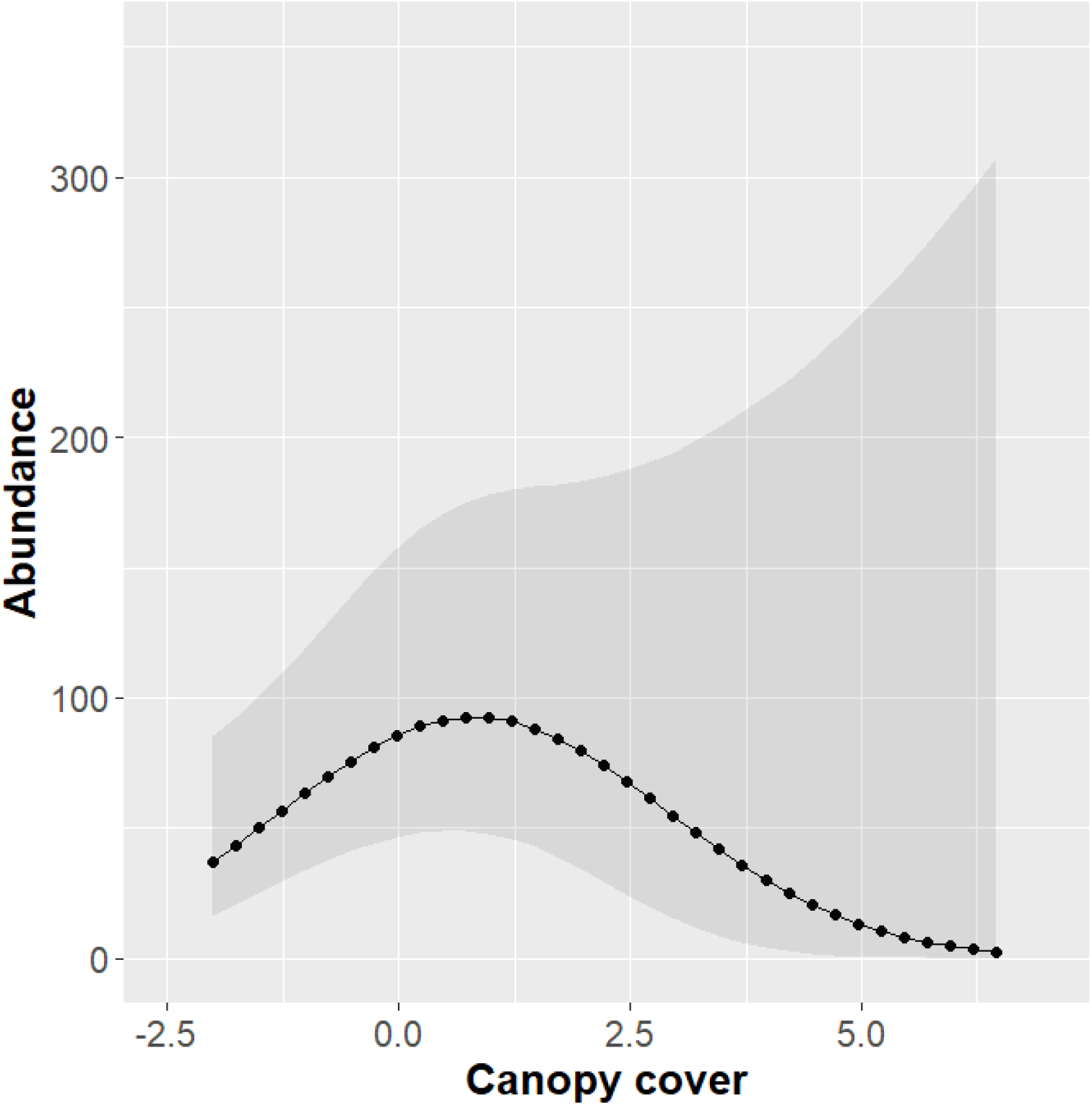
The effect of scaled canopy openness on the abundance of *Adelophryne nordestina* adult males. The shaded area represents the 95% confidence interval

## Discussion

Our results showed that all environmental variables we considered here were a good proxy for the abundance of calling males of *A. nordestina* in PNJVS. Variables such as those considered in this study - leaf litter depth, temperature, rainfall, humidity, and canopy coverage - as well as others, including photoperiod, barometric pressure, and wind speed, affect the calling activity and, consequently, the detectability of certain anuran species (Hatano et al. 2002; Cook et al. 2011; Ospina et al. 2013; Steen et al. 2013; Oseen and Wassersug 2002; Schalk and Saenz 2016; Bonnefond et al. 2019; Ceron et al. 2020). However, environmental variables and the nature of their influence on reproductive activity vary considerably across species or even be opposite among different species within the same community (Schalk & Saenz 2016; Ceron et al. 2020) or in the same species across different seasons, as observed in *Pseudacris crucifer* by Oseen and Wassersug (2002). In addition, there are species in which no influence of environmental variables was found at all (e.g. Dixo & Martins 2008; Ernst & Roddel 2008).

For terrestrial frogs, such as *A. nordestina*, leaf litter depth and humidity are particularly important, since they use the leaf litter as a refuge, calling site, and breeding site (Hoogmoed & Lescure 1984; Hoogmoed et al. 1994; MacCulloch et al. 2008; Lourenço-de-Moraes et al. 2021). Leaf litter depth and humidity were already observed to influence the abundance of leaf litter frogs, including miniaturized and direct development species (Fauth et al. 1989; Van-Sluys et al. 2007). Our results indicate that the species shows a preference for slightly above-average canopy openness, which decreases as the openness approaches a maximum of 35%. The mean canopy openness across all 100 sampled sites was 14.33%, ranging from 4.66% to 35.74%, indicating that, in general, the sampled environments had low light incidence. All these factors together with adequate precipitation contribute to forming a suitable microclimate for the calling activity of *A. nordestina* in the leaf litter of the PNJVS forest. The Abacate, Velha Joana, and Toyô transects, which exhibited the highest abundances and detectability of *A. nordestina*, extended from the forest edge into the interior and were located closer to small streams (see Fig. 1 and 2). These factors likely provide a more humid leaf litter environment, making it more favorable for the species. In contrast, the Podocarpus transect, which recorded the lowest abundance and detectability, extended less into the forest interior and was farther from the streams, characteristics that may have contributed to less suitable conditions for the species due to lower humidity availability.

Within the genus *Adelophryne*, only *A. maranguapensis* has been extensively studied regarding its reproductive biology. This species reproduces in bromeliad phytotelms and exhibits parental care by females (Cassiano-Lima et al. 2011, 2022). Other *Adelophryne* species have only brief mentions regarding their reproductive biology in their descriptions, such as *A. mucronata*, which has been observed in association with terrestrial bromeliads (Lourenço-de-Moraes et al. 2012), and *A. adiastola* (Hoogmoed & Lescure, 1984), *A. glandulata* (Lourenço-de-Moraes et al. 2014), and *A. michellin* (Lourenço-de-Moraes et al. 2018), in which two to three eggs were observed through the females’ skin, suggesting direct development, as proposed by the authors. The terrestrial bromeliads observed in the habitats of *A. nordestina* in PNJVS present narrow leaves, a morphology that does not favor the formation of phytotelmata within them. Therefore, they do not appear to provide a suitable environment for reproduction. We also observed females carrying two to three large eggs throughout all sampling months, with a significant number of juveniles appearing from June onwards. Since the rainy season begins in May, we hypothesize that the species reproduces on the forest floor and requires consistent soil moisture for reproduction. However, further studies on the reproductive biology of the species are necessary to test and confirm this hypothesis.

According to our model, the mean abundance of male *Adelophryne* among sites was 85.9 individuals. Abundance estimates are directly influenced by the species detection probability, and low estimates of detection, such as observed here (mean detection probability 0.03) can bias high abundance estimates. Such abundance estimates, provided by the Negative Binomial model, were four to five times higher than the Poisson model. The Negative Binomial model was already reported as predicting unrealistic high abundance estimates and low detection probabilities before (Kéry 2005, Kéry & Royle 2016). Additionally, previous authors have already cautioned that the Negative Binomial may not represent the best choice to accommodate overdispersion when estimating abundance for N-Mixture models, and Poisson distributions may provide more ecologically realistic estimates (Joseph et al. 2009). However, the full model with Negative Binomial error distribution was favored in relation to the Poisson and the Zero-Inflated Poisson, showing the best fit to our data and posing a conflict between good fit and bad prediction. In this sense, we see our estimates of abundance and detection probability with caution, despite the fact that using a hierarchical modeling approach allowed us to separate the ecological information in the data from inherent sampling errors common in ecological field data.

Overall, we observed that the presence of *A. nordestina* appears to be conditioned by a set of environmental characteristics that shape a suitable habitat for the species within the forest, primarily associated with mountain streams - a hypothesis that should be tested in future studies. Unfortunately, the Brazilian Atlantic Forest is an extremely fragmented environment threatened by anthropogenic pressure, making the existence of conservation units such as PNJVS essential for protecting sensitive species with such specific microhabitat preferences as *A. nordestina*.

Studies addressing the population dynamics of miniaturized frogs remain scarce, not only in Atlantic Forest environments, likely due to challenges in detection, limited distribution, and the difficulty of accessing the locations where these species occur. In this regard, the present study provides an important contribution to understanding how environmental variables may influence the population dynamics and calling activities of these miniaturized frogs, which occupy a unique niche within the Atlantic Forest.

## Acknowledgements

We would like to thank the management of Professor João Vasconcelos Sobrinho Natural Park for allowing and supporting the research, Adriano Andrade for logistical support and assistance in the field during all expeditions, and Ademário Sousa for logistical support. We are grateful to Henrique Andrade for training and help in the pilot fields and to Iza Vilella, Maria Laura Santos, Breno Rafael, Caio Pimentel, Igor Gonçalves, Alison Soares, Will Lins, Carlos Rodrigues, Luana Lyra, Stéphanie Paiva, Carlos Fernandes, Mauro Monteiro, and Edson Silva for helping with the fieldwork. We are also grateful to Pedro Ivo Simões and Alexandre P. de Almeida for their contributions to this paper. This work is the result of LCM’s master’s research, which was supported by Coordenação de Aperfeiçoamento de Pessoal de Nível Superior (Capes) with the scholarship Process: 88887.670698/2022-00 and a grant from the Programa de Apoio à Pós-Graduação (PROAP). KCA was supported by Conselho Nacional de Desenvolvimento Científico e Tecnológico (CNPq) and Fundação de Amparo à Pesquisa do Estado do Piauí (FAPEPI) (Process: 150013/2023-0). IJR was supported by Programa de Desenvolvimento Científico e Tecnológico Regional – PDCTR (CNPq/Funcap Edital 03/2021, DCT-0182-00049.01.00/21 and 04863348/2022) for fellowship (PDCTR 301304/2022-0). This work was carried out under license Sisbio: 80239-1, granted by the Chico Mendes Institute for Biodiversity Conservation - ICMBio. Ethics approval was not required for this study according to Brazilian legislation (Instrução Normativa IBAMA nº 154/2007), as no animals were collected or handled.

